# H3K27me3 is a determinant of chemotolerance in triple-negative breast cancer

**DOI:** 10.1101/2021.01.04.423386

**Authors:** Justine Marsolier, Pacôme Prompsy, Adeline Durand, Anne-Marie Lyne, Camille Landragin, Amandine Trouchet, Sabrina Tenreira Bento, Almut Eisele, Sophie Foulon, Léa Baudre, Kevin Grosselin, Mylène Bohec, Sylvain Baulande, Ahmed Dahmani, Laura Sourd, Eric Letouzé, Elisabetta Marangoni, Leïla Perié, Céline Vallot

**Author notes:** These authors contributed equally to this work (Adeline Durand, Anne-Marie Lyne). Broad Institute of MIT and Harvard, Cambridge MA, USA. co-last authors (Leïla Perié, Céline Vallot).

## Abstract

Triple-negative breast cancer is associated with the worst prognosis and the highest risk of recurrence among all breast cancer subtypes^1^. Residual disease, formed by cancer cells persistent to chemotherapy, remains one of the major clinical challenges towards full cure^2,3^. There is now consensus that non-genetic processes contribute to chemoresistance in various tumor types, notably through the initial emergence of a reversible chemotolerant state^4–6^. Understanding non-genetic tumor evolution stands now as a prerequisite for the design of relevant combinatorial approaches to delay recurrence. Here we show that the repressive histone mark H3K27me3 is a determinant of cell fate under chemotherapy exposure, monitoring epigenomes, transcriptomes and lineage with single-cell resolution. We identify a reservoir of persister basal cells with EMT markers and activated TGF-β pathway leading to multiple chemoresistance phenotypes. We demonstrate that, in unchallenged cells, H3K27 methylation is a lock to the expression program of persister cells. Promoters are primed with both H3K4me3 and H3K27me3, and removing H3K27me3 is sufficient for their transcriptional activation. Leveraging lineage barcoding, we show that depleting H3K27me3 alters tumor cell fate under chemotherapy insult – a wider variety of tumor cells tolerate chemotherapy. Our results highlight how chromatin landscapes shape the potential of unchallenged cancer cells to respond to therapeutic stress.

---

Emergence of resistance phenotypes from initially responding or partially-responding tumors has been modeled as a multi-step process in cancer^7^. Initially, post drug insult, only a pool of persister cells - also called drug-tolerant persister cells (DTPs) - manage to tolerate the cancer treatment and survive^8^. These cells constitute a reservoir of cells from which drug-resistant cells, actively growing under cancer treatment, will ultimately emerge^8–10^. In triple-negative breast cancer (TNBC), both genetic and transcriptomic mechanisms have been proposed to drive cancer evolution towards chemoresistance, through combined selective and adaptive modes of evolution^11^. The recent identification of a multi-clonal reversible drug-tolerant state post neo-adjuvant chemotherapy in patient-derived xenografts (PDX)^6^ suggested that the earliest steps of chemoresistance in TNBC are not driven by genetic alterations, but rather by non-genetic plasticity in multiple cancer cells. Similarly, in other cancer types, persister states have been identified solely by changes in transcriptomic and epigenomic features in response to targeted therapies or chemotherapies^12–15^.

Genetic history of many cancer types has been extensively modelled thanks to both bulk and single-cell approaches^11,16^. In contrast, little is known about the epigenomic heterogeneity and dynamics of acquisition of epigenetic alterations. While recent studies have focused on the evolution of DNA methylation^17,18^-among the most stable epigenetic locks to gene expression, contribution to tumor evolution of more versatile epigenetic modifications, has remained poorly understood. Single-cell methods to map repressive and permissive histone modifications, key players of cellular plasticity, have emerged only recently^19,20^, enabling the study of epigenomic diversity within biological systems. Here, combining single-cell transcriptomics and epigenomics with lineage barcoding, we show that the distribution of H3K27me3 – trimethylation at lysine 27 of histone H3 - is a key determinant of cell fate upon chemotherapy exposure in TNBC, shaping expression programs and cell potential to tolerate chemotherapy.

Resistance to adjuvant chemotherapy for TNBC patients, post-surgery, cannot be easily studied as biopsies are not routinely performed when the disease progresses. To circumvent these limitations, we modelled, *in vivo* and *in vitro*, phenotypes of drug-response observed in patients. *In vivo*, we treated a patient-derived xenograft (PDX) model, established from a patient with residual TNBC^19,21^, with Capecitabine, the standard of care for residual breast tumors. After the first round of chemotherapy treatment, mice displayed a pathological complete response (pCR), but tumors eventually recurred (‘recurrent’) and mice were treated again with chemotherapy, to which tumors responded to various extents (‘responder’ or ‘resistant’) (Fig. 1a and Extended Fig. 1a-c). These recurrent tumors potentially arose from persister cells, surviving initial chemotherapy treatment^9^. We isolated ‘persister’ cells by pooling the fat pad from 6 mice with pCR (Extended Fig. 1d-f). We also generated ‘*residual*’ tumors (n=2) to phenocopy a clinical situation of partial response (Fig. 1a and Extended Fig. 1a and 1d) by administering a moderate dose of Capecitabine (270 mg/kg, half the usual dose).

**Figure 1:**
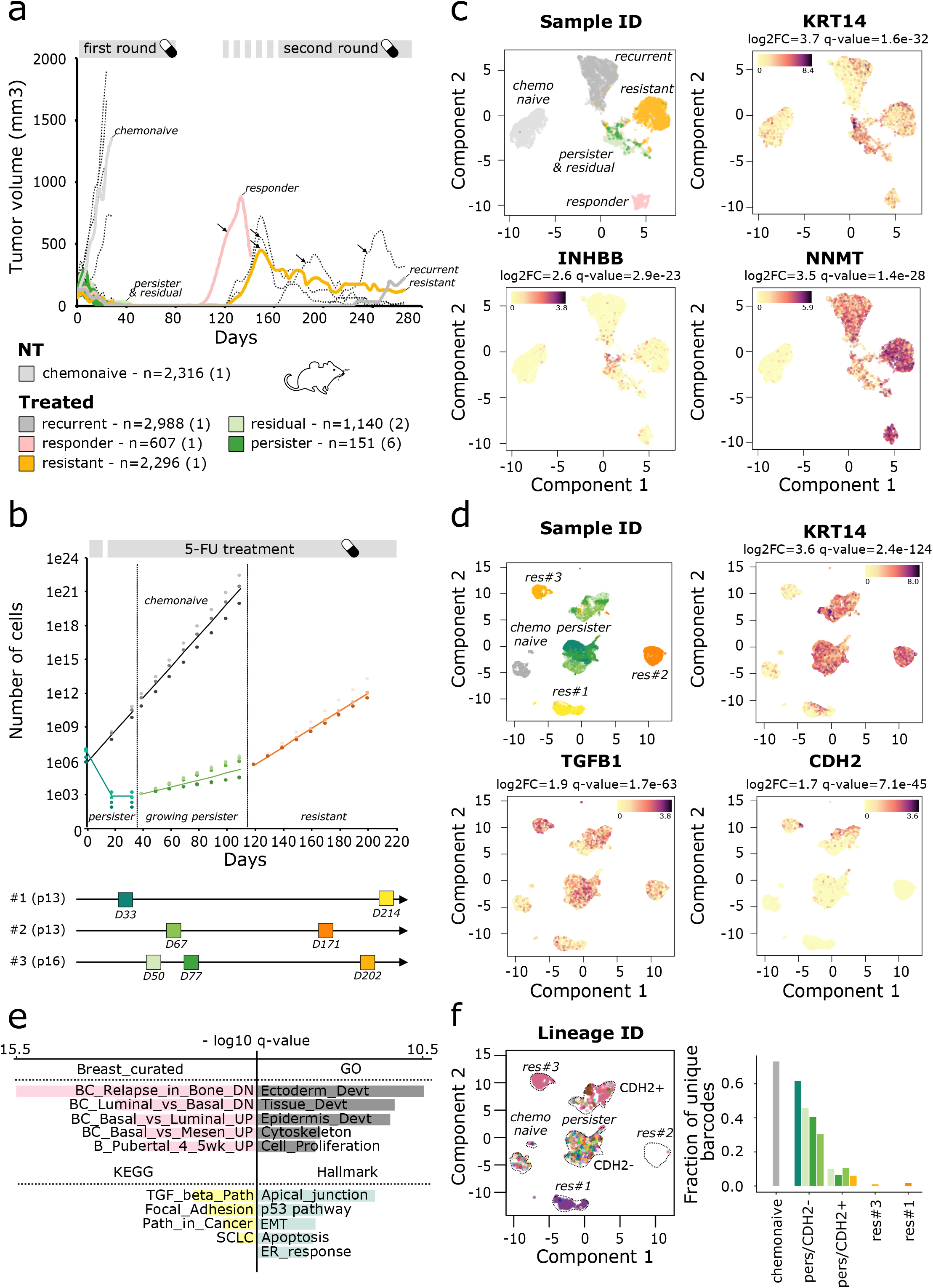
Identification of a pool of basal persister cells in TNBC *in vivo* and *in vitro*. **a.**(Up) Graph of the relative tumor volumes (RTV, mm3) over time (days). Colored growth curves correspond to tumors which have been further studied by scRNA-seq. Black arrows indicate the start of the second round of Capecitabine treatment for the corresponding mice. (Down) Phenotypes and cell numbers are indicated, with the number of mice used to collect samples in brackets. **b.**(Up) Graph representation of the cell proliferation of triple negative breast cancer cell line MDA-MB-468 (MM468) treated with 5-FU (green for persister cells, and orange lines for resistant cells) or with DMSO (chemonaive - grey lines). Each dot corresponds to an independent experiment. (Down) Schematic view of the experimental design. Experience replicate numbers and passages of the cells at D0 are indicated. **c.** UMAP representation of PDX scRNA-seq datasets, colored according to sample of origin (first panel) or gene expression signal for differentially expressed genes between persister cells and chemonaive tumor cells (remaining panels), log2FC and adjusted p-values are indicated above the graph. **d.**UMAP representation of MDA-MB-468 cells scRNA-seq datasets, colored according to the sample ID (first panel) or gene expression signal for differentially expressed genes between persister cells and chemonaive cells (*KRT14* and *TGFB1* panels) or for a differentially expressed gene between the two persisters clusters, *i.e.* clusters C4 vs C2 (*CDH2* panel). Chemonaive population (in grey) corresponds to DMSO-D0-#1. **e.** Barplot displaying the top 5 pathways activated in persister cells both *in vivo* and *in vitro* from MSigDB c2_curated Breast, c2_KEGG, c7_Hallmark and c5_GO annotations. x-axis corresponds to −log10 adjusted p-values for PDX. **f.** (Left) UMAP representation of scRNA-seq as in 1d, cells are colored according to lineage barcode. (Right) Histogram of the lineage barcodes diversity detected in the scRNA-seq data within Louvain partition clusters, and across samples. Colors correspond to sample ID as in 1d.

*In vitro*, we treated an initially chemosensitive TNBC cell line (MDA-MB-468), with the pro-drug of Capecitabine, Fluorouracil (5-FU)^22^, as Capecitabine is not metabolized *in vitro*. We drove independently three pools of cells to chemoresistance with prolonged 5-FU treatment (>15 weeks). After 3 weeks, only few cells survived drug insult (0.01% of the initial population) and started proliferating again under chemotherapy after 10-15 days (Fig. 1b and Extended Fig. 2a-b). Over 15 weeks, populations of resistant cells emerged, with an IC50 to 5-FU over 4-fold higher than chemonaive population and doubling times comparable to chemonaive cells (Extended Fig. 2c). To characterize transcriptomic evolution of chemonaive cells towards chemotolerance and subsequently chemoresistance, we performed single-cell RNA-seq (scRNA-seq) in both cell lines and PDX models (Fig. 1c-d, Extended Fig. 1g-i, 2d-f). *In vivo*, scRNA-seq was mandatory to identify the rare human persister cells among the vast majority of stromal mouse cells. Out of the fat pad, we isolated n~3,480 persister cells, an average of 580 cells per mouse. Both *in vivo* and *in vitro* models, diverse cell populations, with distinct expression programs (Fig.1c-d), originated from the pool of persister cells, which recurrently activated the same set of pathways compared to chemonaive cells (Fig. 1c-e, Extended Fig. 1g-i, Extended Fig. 2d-g). Originating from *KRT5*-expressing cancer cells, persister cells recurrently activated sets of genes further establishing basal cell identity (Extended Fig. 1h-j), such as *KRT14* (Fig. 1c-d). In addition, persister cells showed an activation of the TGF-β pathway with the expression of multiple players including ligands and receptors: *in vivo - TGFB2, INHBA, INHBB, TGFB2R and TGFB3R* (Fig.1c, Extended Fig.1h-i) - and *in vitro* - *TGFB1, INHBA* and *TGFBR1* (Figure 1d, Extended Fig. 2f). Compared to chemonaive cells, persister cells also showed an activation of genes associated to the Epithelial–to-Mesenchymal Transition (EMT, Fig.1c-d, Extended Fig. 1h-i, 2e-f), such as *FOXQ1,* a transcription factor previously identified to drive EMT^23,24^ in cancer, and *NNMT*, characteristic of the metabolic changes that accompany EMT^25–27^. TGF-β and EMT associated-genes have been shown as markers of residual TNBC^28,29^ and were proposed as potential drivers of chemoresistance in lung^30^, pancreatic^31^, breast^32^, and colorectal cancers^33,34^. *In vivo*, we showed that persister and residual tumor cells actually clustered together (cluster C1) and shared a common expression program, suggesting similar mechanisms of chemotolerance independent of the residual burden (Extended Fig. 1g). Yet persister cells displayed a decreased proliferation rate as attested by a higher number of cells in G0/G1 (Extended Fig. 1f and Extended Fig. 2b), in line with previous reports^12,14,35^. *In vitro*, we identified two clusters of persister cells (clusters C2 and C4), that differ by their expression of additional EMT markers such as *CDH2* (Fig. 1d) and *TWIST1.* Early individual persisters (day 33) solely belonged to cluster C2/CDH2- whereas growing persisters could either belong to C2/CDH2− or C4/CDH2+ (Fig.1d and Extended Fig. 2d). Overall, we identified both *in vivo* and *in vitro* a reservoir of persister basal cells with EMT markers and activated TGF-β pathway evolving to multiple resistant phenotypes (Fig. 1c-d). We pinpoint TGF-β and EMT pathway activation as the earliest common molecular events at the onset of chemoresistance in TNBC, defining a common Achilles’ heel, to target chemoresistant cells before they phenotypically diversify.

To follow clonal evolution under therapeutic stress, we had initially introduced unique genetic barcodes in chemonaive MDA-MB468 cells prior to our experiments (Extended Fig. 3a). We leveraged our previous barcoding method^36^ to allow robust detection of barcodes in scRNA-seq data, as shown by the high fraction of cells with a lineage barcode (Extended Fig. 3b). In addition, we verified that barcodes frequencies detected in scRNA-seq data correlated with those detected in bulk, confirming the sensitivity of barcode detection in scRNA-seq data (Extended Fig. 3c). Then comparing barcode diversity at the persister stage, we observed that 5-FU and DMSO-treated cells display equivalent lineage diversity (Extended Fig. 3d-e), showing that the persister state is multi-clonal. To test if the lineages that persist are a random draw of the chemonaive population, we compared barcode frequencies between the starting population and the 5FU or DMSO-treated cells. As cells after barcode tagging and before chemotherapy treatment are growing, resulting in several cells with the same barcodes, if surviving cells had no particular predisposition then they should resemble a random draw of the chemonaive barcoded cells (day 0). This is what we indeed observed for DMSO-treated cells when compared to random drawing (Extended Fig. 3f-g). However, barcode frequencies of the 5-FU treated cell deviate from the random scenario (Extended Fig. 3f-g), indicating that some lineages present in the chemonaive population have a predisposition to tolerate the treatment. This was further confirmed by the comparison of barcode frequencies across persister states from independent experiments showing that independent persister populations shared lineage barcodes (Extended Fig. 3h). To monitor clonal evolution within the different subgroups of persister cells, we compared the barcode diversity within expression clusters (Fig. 1f). We found that lineage diversity decreases over the course of treatment, eventually leading to few clones dominating the chemoresistant cluster (Fig. 1f). The diversity in the CDH2+ persister cluster was lower than in the CDH2-persister cluster (Fig. 1f), suggesting that only rare persister cells switched to the CDH2+ persister state. By combining detection of lineage barcode and expression programs at single-cell resolution, we demonstrated that a small pre-disposed subset of cells tolerate chemotherapy, progressively transiting from a CDH2-to a CDH2+ persister state, and eventually leading fewer cells to resist.

To hamper this chemo-driven clonal evolution of cancer cells, we next investigated the molecular basis of such rapid phenotypic evolution. Using whole-exome sequencing (Extended Fig. 4a), we first analyzed mutations, copy-number alterations (CNA) and related mutational signatures acquired by persister and resistant cell populations since the onset of 5-FU treatment (‘chemonaive D0’ population). We could not identify any recurrent mutations across experiments (Extended Fig. 4b), or any CNA (amplifications or homozygous deletions) or mutations affecting known driver genes of breast cancer in any population^16^. Only a minor fraction of mutations found in persister cells were attributed to 5-FU (mutational signature 17^37^) in comparison to resistant cell populations where over 50% of acquired mutations are associated to 5-FU exposure (p<10^−10^, Extended Fig.4c), indicating that chemo-related mutations are acquired over a timeframe that is not compatible with the rapid phenotypic evolution seen in persister cells. Finally, computing cancer cell fractions for each mutation, we confirmed that persister populations are extensively multi-clonal compared to chemonaive cells (Extended Fig. 4d), in line with the lineage barcoding results.

We next investigated changes in epigenomes during chemotherapy treatment. Using single-cell profiling (scChIP-seq) of the repressive H3K27me3 epigenomic mark, we observed that H3K27me3 epigenomes faithfully captured the evolution of cell states with chemotherapy (Fig. 2a, Extended Fig. 5a-c). Persister cells shared a common H3K27me3 epigenome (Fig. 2a-c, cluster C1), in contrast to resistant cells split in clusters C1 and C3. In comparison to chemonaive cells, cells from cluster C1 showed recurrent redistribution of H3K27me3 methylation, the highest changes (|log2FC|>2) occurring specifically at transcription start sites (TSS) and gene bodies (Fig. 2d) and corresponding to a loss of H3K27me3 enrichment (75 regions with log2FC<-2, and 2 regions with log2FC>2). This depletion correlated with the highest changes in gene expression observed by scRNA-seq (Fig. 2e-f and Extended Fig. 5d) and was associated to the transcriptional de-repression of 14% of *persister* genes (Fig. 2g) – genes overexpressed in persister versus chemonaive cells. In the chemonaive cells, two epigenomic subclones were recurrently identified indicated an epigenomic heterogeneity in this population (n=3 experiments, C2 & C4, Fig. 2b and Extended Fig. 5e-f). In contrast to cells from cluster C4 (median correlation score r=-0.34, no cells over r=0.20), a fraction of cells within cluster C2 shared epigenomic similarities with cells from C1 (49/381 cells over r=0.20, Fig. 2c and Extended Fig. 5g-h), but remained discernible from the pool of persister cells (no cells from C2 over median r of C1, see Methods). This suggests that cells from C2 could fuel the persister population when exposed to chemotherapy, with the need of chemo-induced chromatin changes to achieve tolerance. In addition to these two epigenomic subclones within the chemonaive cells, we also detected rare cells with a persister epigenomic signature, in only one of our three experiments (60/976 chemonaive-D60-#1 cells in cluster C1 – Extended Fig. 5f), suggesting that spontaneous transition to H3K27me3 persister state rarely occurs in the absence of chemotherapy.

**Figure 2:**
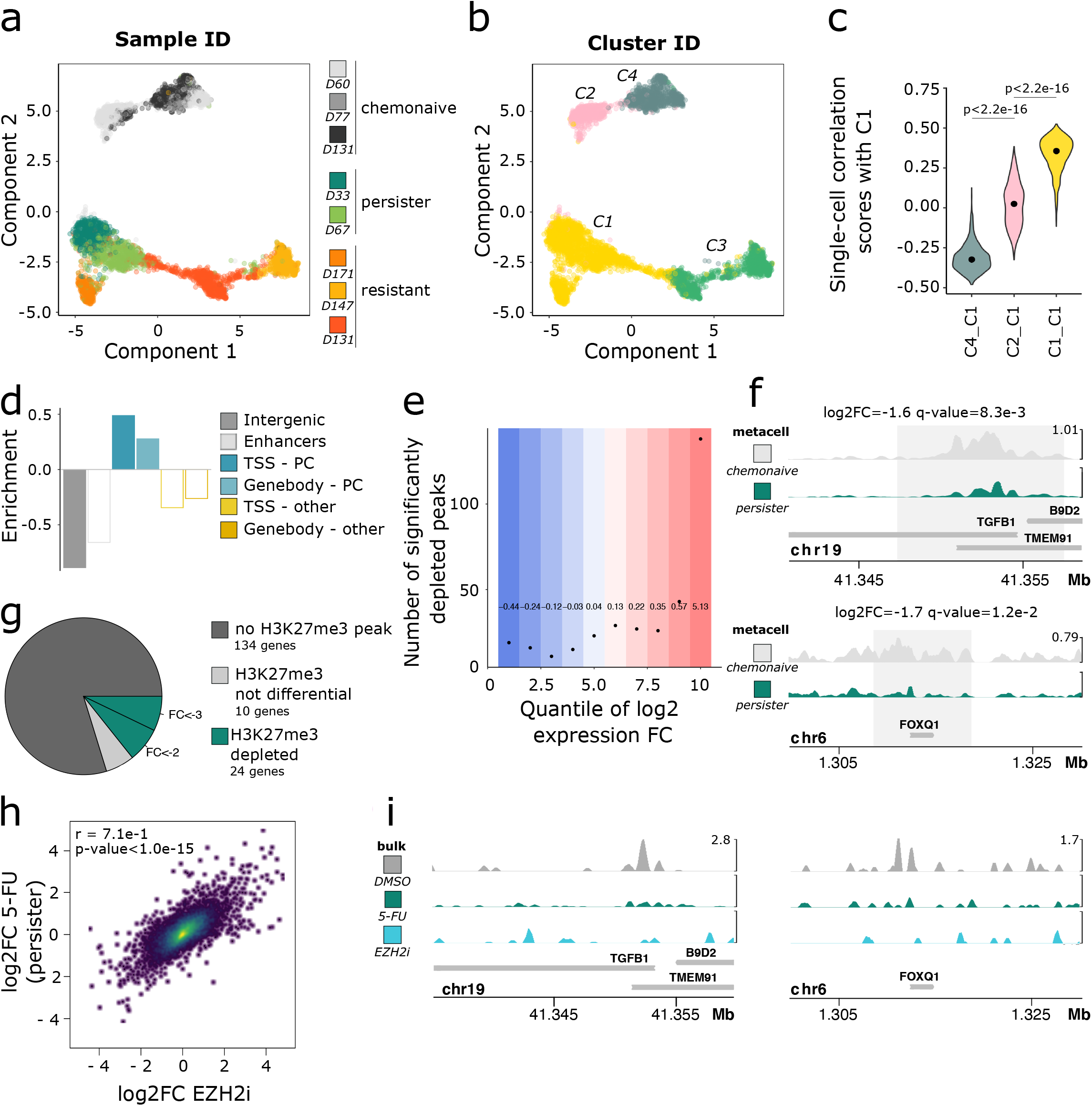
H3K27me3 represses the persister expression program in chemonaive cells. **a.** UMAP representation of scChIP-seq H3K27me3 datasets, cells are colored according to the sample of origin. Chemonaive samples correspond to DMSO-treated cells, persister and resistant samples correspond to 5-FU-treated cells, days of treatment are indicated. **b.** Same as in a. with cells colored according to epigenomic clusters. **c.** Graph representation of the cell to cell inter-correlation between clusters C1, C2 or C4 and the cluster C1. **d.**Genomic association between H3K27me3 peaks and gene annotation. Full bars indicate adjusted p-value<1.0 10^-2^. Empty bars indicate non-significant adjusted p-values. “PC” indicates protein coding gene **e.** Repartition of H3K27me3 depleted peaks within re-expression quantiles in persister cells. **f.** Cumulative scH3K27me3 profiles over *TGFB1* and *FOXQ1* in chemonaive and persister cells (D33). Log2FC and adjusted p-value correspond to differential analysis comparison of cells from cluster C1 versus clusters C2 + C4. **g.**Pie chart displaying the fraction of persister genes potentially regulated by H3K27me3 in MM468 cells. **h.**Dot plot representing log2 expression fold-change induced by 5-FU or EZH2i at D33 versus D0. Correlation scores and associated p-value are indicated. **i.** Bulk H3K27me3 chromatin profiles for *TGFB1* and *FOXQ1* in cells treated with DMSO, 5-FU or EZH2i at D33.

To test whether H3K27me3 enrichment was the lock to the persister expression program in chemonaive cells, we treated cancer cells with the EZH2 inhibitor (EZH2i) UNC1999^38^, to deplete H3K27me3 from cells (Extended Fig. 5i), in the absence of chemotherapy. EZH2i treatment phenocopied persister state to chemotherapy as expression fold-change induced by EZH2i were specifically correlated to those induced by chemotherapy exposure at early time points (Fig. 2h, Extended Fig. 5j, r=0.71 versus r=0.31 with changes in resistant cells). Furthermore, we observed that EZH2i was sufficient to lead to the activation of 62% of H3K27me3-enriched *persister* genes (15/24 genes), suggesting that H3K27me3 was the sole lock to their activation (Fig. 2i and Extended Fig. 5k-l). EZH2i was also sufficient to lead to the over-expression of 61% of *persister* genes independently of any H3K27me3 enrichment in chemonaive cells (86/144 genes), such as *KRT14*, suggesting that these genes might be targets of H3K27me3-regulated *persister* genes.

As we observed H3K27me3 changes precisely at TSS, we further explored the evolution of chromatin modifications at TSS, focusing on H3K4me3 - trimethylation at lysine 4 of histone H3 - a permissive histone mark shown to accumulate over TSS with active transcription. In contrast to single-cell H3K27me3 epigenomes which were sufficient to separate cell states along treatment (Fig 2a), individual H3K4me3 epigenomes of chemonaive and persister cells were indiscernible (Fig. 3a and Extended Fig. 6a-b). Comparing H3K4me3 single-cell tracks of chemonaive and persister cells (Fig. 3b and Extended Fig. 6c), we observed sparse H3K4me3 enrichment at the TSS of *persister* genes already in chemonaive cells, compared to house-keeping genes (Extended Fig. 6d). In individual persister cells, H3K4me3 enrichment was significantly more synchronous at these TSS (Extended Fig. 6e, p=9.2 10^−3^) than in chemonaive cells. Altogether, H3K4me3 and H3K27me3 could either accumulate on the same TSS but in different chemonaive cells, or H3K4me3 could accumulate together with H3K27me3 in a subset of cells.

**Figure 3:**
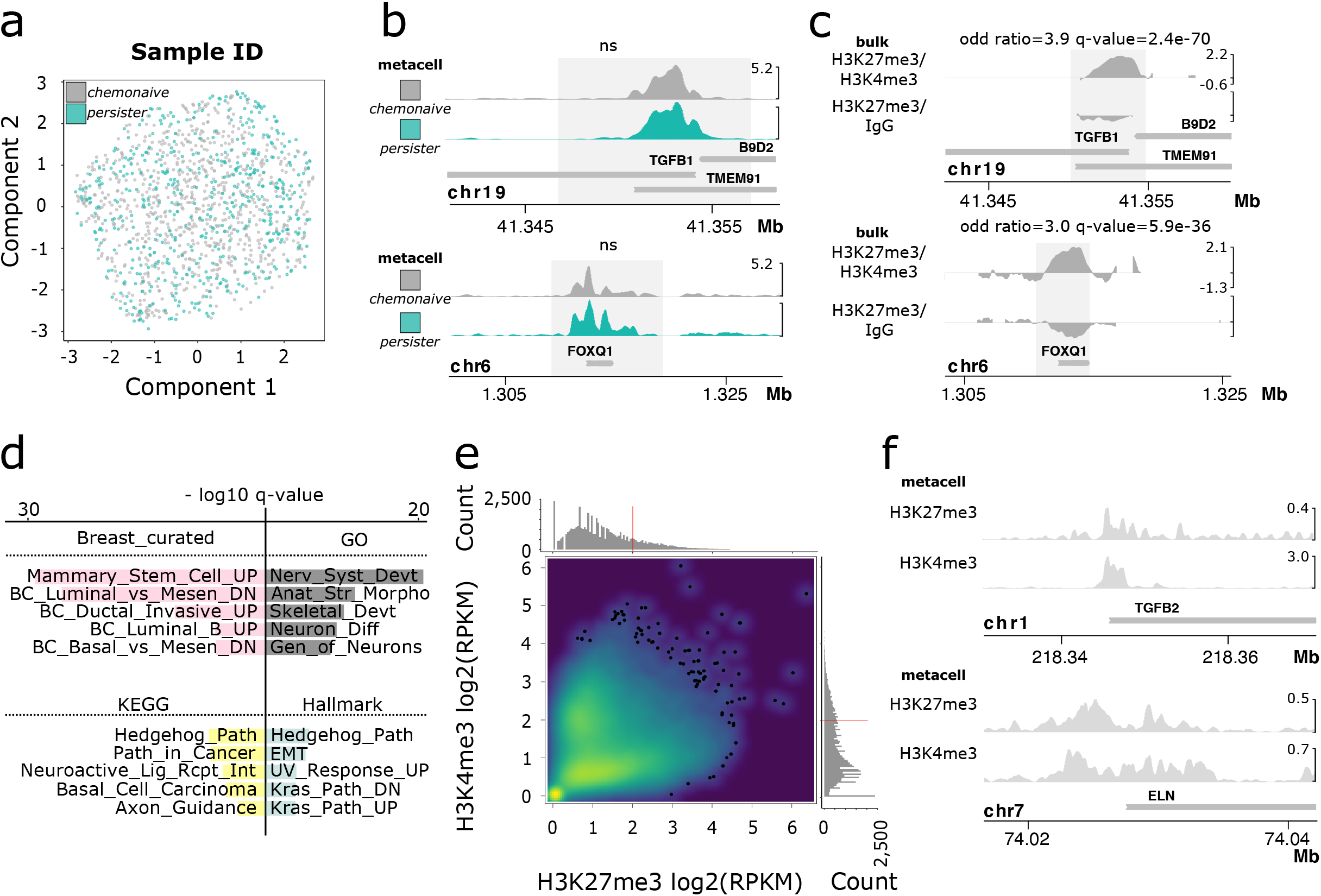
Epigenomes of chemonaive cells are primed with co-accumulation of H3K27me3 and H3K4me3. **a.** UMAP representation of scChIP-seq H3K4me3 datasets, cells are colored according to the sample of origin. **b.** Cumulative scH3K4me3 enrichment profiles over *TGFB1* and *FOXQ1* in chemonaive cells (D0) and persister cells (D60). **c.** Bulk H3K27me3->H3K4me3 and H3K27me3->IgG chromatin profiles of *TGFB1* and *FOXQ1* in the chemonaive population (D0). Enrichment tracks show enrichment over IgG control with associated odd ratio and adjusted p-value. **d.** Barplot displaying the top 5 pathways (as in Fig. 1e) enriched in genes detected with combined H3K27me3/H3K4me3 in the chemonaive MDA-MB-468 cell population (D0). **e.** Density plot representing cumulative scH3K27me3 and scH3K4me3 log2 enrichment at TSS in two independent cell populations within chemonaive PDX. **f.** Cumulative scH3K27me3 and scH3K4me3 profiles over *TGFB2* and *ELN* in the chemonaive PDX tumor.

To test whether H3K4me3 could co-exist with H3K27me3 in the same individual cells prior to chemotherapy exposure, we performed successive immunoprecipitation of H3K27me3 with H3K4me3 or H3K27me3 with isotype control (IgG). We detected n= 1,547 transcription start sites significantly enriched in DNA immunoprecipitated with both H3K27me3 and H3K4me3, compared to the control (H3K27me3/IgG) precipitated fraction (peak-ratio>0.15, q-value<1.0 10^−3^, Extended Fig. 6g). We found that bivalent chromatin in chemonaive cells (D0) was detected at genes associated to basal and EMT pathways, as well as various developmental pathways (e.g Hedgehog pathway) (Fig. 3c-d and Extended Fig. 6f), and at the ligand of the TGF-β pathway, TGF-β1 (Fig. 3c). The majority of K27-regulated *persister* genes (18 out of 24) were found in a bivalent chromatin configuration in the chemonaive cell population (Fig. 3c, Extended Fig. 6f). *In vivo*, we observed enrichment of H3K27me3 and H3K4me3 modifications measured independently at n=1,370 TSS, particularly at EMT-associated genes and TGF-β2 (Fig. 3e-f and Extended Fig. 6h-i), corroborating that *persister* genes are in a bivalent configuration in chemonaive cells.

To further validate that H3K27me3 distribution regulates the emergence of persister cells, we next modulated H3K27me3 distribution simultaneously to chemotherapy. We used both the EZH2 inhibitor (EZH2i - UNC1999^38^), and a KDM6A/B inhibitor (KDM6A/Bi - GSK-J4^39^) to prevent demethylation of H3K27me3 residues. Co-treating cells with one of these modulators together with 5-FU, we observed opposite modulation of the ability of cells to tolerate the chemotherapy. EZH2i increased the number of persister cells, whereas KDM6A/Bi led to a decrease in the number of persisters at day 21 (Fig. 4a-b). At day 60, KDM6A/Bi further completely abolished the growth of persister cells under 5-FU, whereas it has no effect on chemonaive cancer cells (Fig. 4c and Extended Fig. 7a). These results were confirmed in a second TNBC cell line HCC38, albeit to a lesser extent for EZH2i (Extended Fig. 7b-c). We then tested how EZH2i affected cell fate of persister cells. Comparing the number of unique barcodes present in the two clusters under chemotherapy pressure with or without EZH2i (Extended Fig. 3a), we observed that co-treating cells with EZH2i and 5-FU increases the number of lineage barcodes in persister cells, and to a large extent in C4/CDH2+ cluster (Fig. 4d-f). These results showed that under EZH2i cancer cells have an increased potential to reach persister state, and a mesenchymal CDH2+ state. Overall, inhibiting the removal of methyl groups at H3K27 prevented cells from reaching chemotolerance and resistance, demonstrating that some cancer cells need to actively demethylate H3K27me3 residues to reach persister state and to proliferate under chemotherapy exposure. Conversely, depleting H3K27me3 from chemonaive cells not only launched a persister-like expression program, but it also enhanced chemotolerance. These results were consistent with a mechanism where persister genes would be repressed by H3K27me3 in chemonaive cells, and primed with stochastic H3K4me3 in a subset of cells – loss of H3K27me3 being the key to activation.

**Figure 4:**
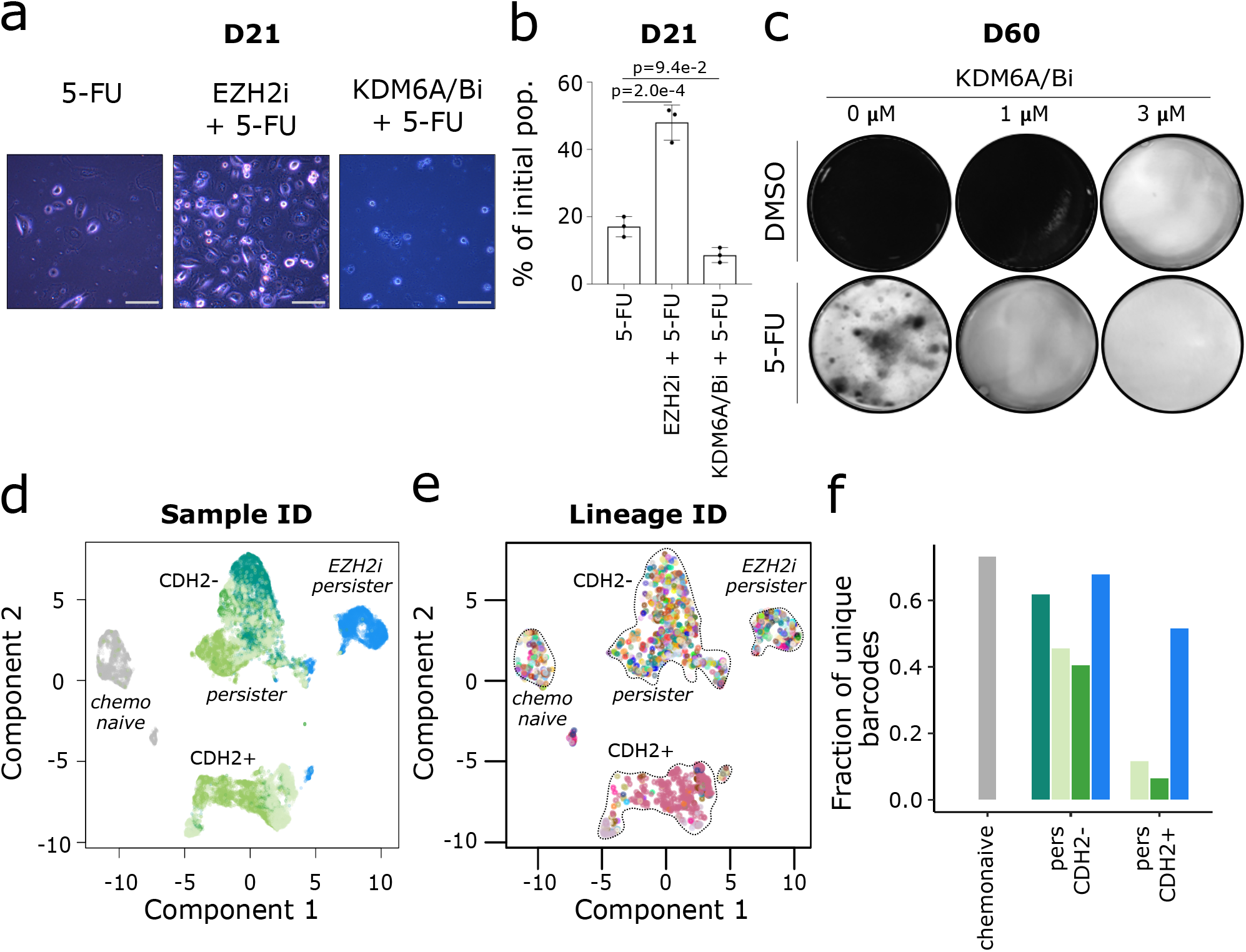
EZH2 inhibition modulates cell fate upon chemotherapy exposure. **a.** Representative pictures of MDA-MB-468 cells treated 21 days with 5-FU alone or in combination with an inhibitor of EZH2 (UNC1999) or an inhibitor of KDM6A/B proteins (GSKJ4). **b.** Histogram representing the number of cells after treatment with 5-FU alone, 5-FU and EZH2i or KDM6i over 21 days, relative to the number of cells at D0. (n=3, Mean ± sd). **c.** Colony forming assay of MDA-MB-468 co-treated with DMSO or 5-FU and indicated concentrations of KDM6A/Bi for 60 days. The data corresponds to 1 of 3 biological replicates. **d.** UMAP representation of scRNA-seq datasets, colored according to the sample ID. **e.** As in 4d. cells colored according to lineage barcode. **f.** Histogram of the diversity of lineage barcodes detected in scRNA-seq data in each louvain partition cluster obtained from the chemonaive population, 5-FU or EZH2i persister cells (D33). Colors correspond to sample ID as in 4d.

In conclusion, our study shows that the transition to persister state in TNBC is dependent on the control of H3K27me3 distribution. We propose that combining chemotherapy with histone demethylase inhibitors at the onset of chemotherapy exposure will decrease the pool of persister cells, and thereby decrease recurrence. We demonstrate a role for both H3K27me3 writer and erasers in regulating the phenotypic switch from chemonaive to chemopersister state, highlighting the instrumental role of repressive histone landscapes as determinants of cell fate. Several studies had started to interrogate which epigenetic modifiers could regulate expression programs of persister or resistant cells, but studying them in isolation^5,12,14,40,41^. We also show that the *persister* expression program is primed in chemonaive cells with a stochastic presence of H3K4me3 but is repressed by H3K27me3. In other words, genes are ready to be activated, H3K27me3 remaining the only lock to activation. Persisters are cells without H3K27me3 or the one releasing the H3K27me3 lock, or a mixture of both phenomena. We observe epigenomic priming of signaling pathways known to participate to drug resistance in TNBC^42^, including Hedgehog, WNT (Extended Fig. 2f), TGF-β and ATP-binding cassette drug transporters pathways (Extended Fig. 2f). Such epigenomic priming is reminiscent of developmental bivalency priming mechanisms^43^ found in stem cells prior to differentiation and appears key for the rapid activation of the genes upon therapeutic stress. Remains to be understood, how only a fraction of bivalent genes are targeted by gene activation upon chemotherapy exposure, whether tolerance to different drugs triggers the activation of the same set of bivalent genes, and whether such mechanisms could be shared across cancer types. Determining the precise addressing mechanisms that target H3K27me3 and H3K4me3 writers and readers to TSS of bivalent genes in cancer cells, will be instrumental in the future to identify dedicated co-factors which could serve as alternative therapeutic targets to restrict the epigenetic plasticity of cancer cells.

## Methods

### PDXs

Female Swiss nude mice were purchased from Charles River Laboratories and maintained under specific-pathogen-free conditions. Mouse care and housing were in accordance with institutional guidelines and the rules of the French Ethics Committee (project authorization no. 02163.02). In this study, we used a xenograft model generated from a residual triple-negative breast cancer post-neoadjuvant chemotherapy (HBCx95) established previously at Curie Institute with informed consent from the patient^44,45^. Five mice were not treated and kept as controls and twenty-seven mice were treated orally with Capecitabine (Xeloda; Roche Laboratories) at a dose of 540 mg/kg, 5 d/week for 6 to 14 weeks. Relative tumor volumes (mm3) were measured as described previously^21^. Latency was defined as the number of days between the observation of a complete response (Tumor size < 10 mm3) after the first round of Capecitabine treatment, and the detection of a recurrent tumor (Tumor size > 10 mm3). Eight mice were sacrificed after the first round of chemotherapy to study residual (2 mice) and persister (6 mice) human tumor cells. Seven mice with recurrent tumors (tumor volume between 200 and 600 mm3) were treated for a second round of Capecitabine. The GraphPad PRISM 9 was used for statistics in Extended Fig.1b. The results represent the mean ± sd and statistical analysis was performed using two-tailed Mann-Whitney test.

Before downstream analysis (scChIP-seq, scRNA-seq), control and Capecitabine treated tumors were digested for 2 h at 37 °C with a cocktail of Collagenase I (Roche, Ref: 11088793001) and Hyaluronidase (Sigma-Aldrich, Ref: H3506). Cells were then individualized at 37°C using a cocktail of 0.25% Trypsin-Versen (Thermo Fisher Scientific, Ref: 15040-033), Dispase II (Sigma-Aldrich, Ref: D4693) and DNase I (Roche, Ref: 11284932001) as described previously^46^. Then, eBioscience red blood cell lysis buffer (Thermo Fisher Scientific, Ref: 00-4333-57) was added to the cell suspension to remove red blood cells. To increase the viability of the final cell suspension, dead cells were removed using the Dead Cell Removal Kit (Miltenyi Biotec, Ref:130-090-101).

### Cell lines, culture conditions and drug treatments

MDA-MB-468 cells were cultured in DMEM (Gibco-BRL, Ref: 11966025), supplemented with 10% heat-inactivated fetal calf serum (Gibco-BRL, Ref: 10270-106). HCC38 cell lines were cultured in RPMI 1640 (Gibco-BRL, Ref: 11875085), supplemented with 10% heat-inactivated fetal calf serum. All cell lines were cultured in a humidified 5% CO2 atmosphere at 37 °C, and were tested as mycoplasma negative. GSKJ4 (KDM6A/B inhibitor, Sigma, Ref: SML0701), GSKJ5 (GSK-J4 inactive isomer, Abcam, Ref: ab144397) and UNC1999 (EZH2 inhibitor, Abcam, Ref: ab146152) were used at indicated concentrations. Cells were treated with 5 μM of 5-FU (Sigma, Ref: F6627) alone or in combination with KDM6A/Bi or EZH2i for indicated days.

### Colony forming assay

TBNC cells were plated in 6 multi-well plates at a density of 200,000 cells per well and treated with the indicated drugs for 60 days (MDA-MB-468) or 50 days (HCC38). Treated plates were monitored for growth using a microscope. At the time of maximum foci formation, colony formation was evaluated after a staining with 0.5% Crystal Violet (Sigma, Ref: C3886).

### Cell proliferation, doubling time and IC50

MDA-MB-468 and HCC38 cells were stained with Trypan Blue (Invitrogen, Ref: T10282) exclusion test, and counted using a Countess automated cell counter (Invitrogen, Ref: C10228) at indicated time of treatment (Fig.4b and Extended Fig.7b). Doubling time (Extended Fig.2c) was calculated using this formula:

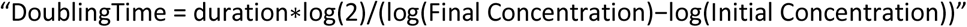

For chemonaive condition and resistant condition, cell numbers were evaluated on cell population during 10 days (n=3). For persister condition, cells were counted manually under the microscope at day 13 and day 30 of treatment. Doubling time of 5-FU growing persister cells was studied from single cell to confluent colony by assaying cell number every 4 days during 27 days (n= 6 single cells) (Figure 1b).

MDA-MB-468 chemonaive and chemoresistant cells were plated in 96 multi-well plates at a density of 10,000 cells per well and treated with increased concentration of 5-FU (1μM to 0.5M) for 72h. Cell cytotoxicity was assayed with XTT kit (Sigma, Ref: 11465015001) and IC50 was calculated as the concentration of 5-FU that is required to obtain 50% of cell viability (Extended Fig. 2c).

The GraphPad PRISM 9 was used for statistics and the results represent the mean ± sd of three independent experiments. Statistical analysis was performed using the Bonferroni test for multiple comparisons between samples (Fig.4b and Extended Fig.2c-right) or one-tailed Mann-Whitney test for the comparison between two conditions (Extended Fig. 2c-left and Extended Fig.7b).

### Western blotting

DMSO- and EZH2i-treated cells (D33-#8) were lysed at 95°C for 10 minutes in Laemmli buffer (50 mM Tris-HCl [pH 6.8], 2% SDS, 5% glycerol, 2 mM DTT, 2.5 mM EDTA, 2.5 mM EGTA, 4 mM Sodium Orthovanadate, 20 mM Sodium Fluoride, protease inhibitors, phosphatase inhibitors) and proteins concentrations were measured using a Pierce BCA protein Assay Kit (Thermo Fisher Scientific, Ref: 23225/23227). 10 μg of proteins were then separated on a 4-15% Mini-PROTEAN TGX Stain-Free Gel (Bio-Rad, Ref: 4568085) at 160V. After transfer, the membrane was exposed to UV light (Bio-Rad, ChemiDoc MP) and the image was further used for total protein quantification. The membrane was then blocked for 1 h at room temperature in PBS pH 7.4 containing 0.1% Tween-20 and 1% milk (Regilait). Incubation anti-H3K27me3 (Dilution: 1:2000, Cell Signaling, Ref: 9733) primary antibodies diluted in PBS pH 7.4, 0.1% Tween-20 were performed at 4°C overnight. Following 2 h incubation at room temperature with an anti-rabbit peroxidase-conjugated secondary antibody (Dilution: 1:10000, Thermo Fisher Scientific, Ref: 31460) diluted in PBS pH 7.4, 0.1% Tween-20, antibody-specific labeling bands were revealed (Bio-Rad, ChemiDoc MP) using a SuperSignal West Pico PLUS Chemiluminescent Substrate (Thermo Fisher Scientific, Ref: 34579).

### Lentivirus packaging and cell transduction

Lentivirus was produced by transfecting the barcode plasmids pRRL-CMV-GFP-BCv2AscI and p8.9-QV and pVSVG into HEK293T cells as previously described^36^. MDA-MB-468 cells from ATCC were infected at passage 11 with lentivirus produced from the barcode library (pRRL-CMV-GFP-BCv2AscI) which includes 18206 different barcodes of 20bp of a random stretch, at a low multiplicity of infection (MOI 0.1) to minimize the number of cells marked by multiple barcodes. Three weeks after transduction, cells were sorted for GFP expression to select cells with barcode insertion, and used for drug treatment.

### Single-cell RNA-seq

For each single cell suspension (DMSO-D0-#1, 5-FU-D33-#1, 5-FU-D214-#1, 5-FU-D67-#2, 5-FU-D171-#2, 5-FU-D50-#3, 5-FU-D77-#3 and 5-FU-D202-#3), approximately 3,000 cells were loaded on a Chromium Single Cell Controller Instrument (Chromium Single Cell 3ʹv3, 10X Genomics, Ref: PN-1000075) according to the manufacturer’s instructions. Samples and libraries were prepared according to the manufacturer’s instructions. Libraries were sequenced on a NovaSeq 6000 (Illumina) in PE8-8-91 with a coverage of 50,000 reads/cell.

### Bulk lineage barcode library preparation and sequencing

Lineage barcodes are recovered by isolating genomic DNA from cells of interest (NucleoSpin Tissue, Mini kit for DNA from cells and tissue, Macherey Nagel, Ref: 740952.50). From the isolated genomic DNA, barcodes are amplified with three nested PCR steps as decribed in^36^. In short, after a first specific PCR for the common region of the lineage barcodes, the amplified material was prepared for sequencing by addition of the illumina sequencing adaptaters and indexing and purification. Sequencing was done in order to obtain 50 reads, on average, per barcoded cell.

### Bulk ChIP-seq

ChIP experiments were performed as previously described^19^ on 3×10^6^ MDA-MB-468 cells (DMSO-D67-#2, DMSO-D77-#3, DMSO-D113-#4, 5-FU-D67-#2, 5-FU-D77-#3, 5-FU-D113-#4) using an anti-H3K27me3 antibody (Cell Signaling Technology, Ref: 9733-C36B11). Sequencing libraries were prepared using the NEBNext Ultra II DNA Library Prep Kit (NEB, Ref: E7645S) according to the manufacturer’s instructions. Libraries were sequenced on a NovaSeq 6000 (Illumina) in SE50 mode.

### Single-cell ChIP-seq

Cells (DMSO-D60-#1, DMSO-D77-#3, DMSO-D131-#5, 5-FU-D33-#1, 5-FU-D67-#2, 5-FU-D171-#2, 5-FU-D147-#3, 5-FU-D131-#6) were labeled by 15 min incubation with 1 μM CFSE (CellTrace CFSE, ThermoFisher Scientific, Ref: C34554). Cells were then resuspended in PBS supplemented with 30% Percoll, 0.1% Pluronic F68, 25 mM Hepes pH 7.4 and 50 mM NaCl. Cell encapsulation, bead encapsulation and 1:1 droplet fusion was performed as previously described^19^. Immunoprecipitation, DNA amplification and library were performed as in^19^. Libraries were sequenced on a NovaSeq 6000 (Illumina) in PE100, with 4 dark cycles on Read 2, with a coverage of 100,000 reads/cell.

### Quantitative chromatin profiling with chromatin indexing

Chromatin isolation, indexing, immunoprecipitation and library preparation was adapted from^47^. Briefly, 50,000 MDA-MB-468 were lysed and digested with MNase for 20min at 37°C in the following buffer: 46mM Tris-HCl pH 7.4, 0.154M NaCl, 0.1% Triton, 0.1% NaDoc, 4.65mM CaCl2, 0.47x Protease Inhibitor Cocktail (Roche, Ref: 11873580001) and 0.09u/uL MNase (Thermo Scientific, Ref: EN0181). Fragmented nucleosomes were then ligated for at least 24h at 16°C to double-stranded barcoded adapters containing 8bp barcodes to combine samples: Pac1-T7-Read2-8bpBarcode-linker-Pac1. Next, 5 indexed chromatin samples (DMSO, 5-FU, UNC, 5-FU + UNC, GSK-J4) were pooled, each containing a different 8-bp barcode, to perform anti-H3K27me3 ChIP (Cell Signaling, Ref: 9733-C36B11) on 250,000 cells in total in each pool. ChIP and DNA amplification was carried out as for scChIP-seq^19^ and a sequencing library was produced for both IP and input pools and sequenced on NovaSeq 6000 (Illumina) in PE100 mode.

### Sequential ChIP-seq

Primary ChIP experiments were performed as described previously^19^ on 30x 10^6^ untreated chemonaive MDA-MB-468 cells using the anti-H3K27me3 antibody. After washes, samples were eluted twice at 37°C for 15 min under agitation in an elution buffer (50mM Tris-Hcl pH8, 5mM EDTA, 20mM DTT, 1% SDS) as in^48^. Samples were diluted 10 times to decrease SDS and DTT concentration. 10% of the eluted chromatin was kept as primary ChIP. Secondary ChIP, re-ChIP, was performed overnight on the rest of the primary immuno-precipitated chromatin using an anti-H3K4me3 antibody (Cell signaling, Ref: #9751) or using an anti-IgG antibody (Cell signaling, Ref: #3900) as a control, to determine the background level of the re-ChIP experiment. After washes, samples were eluted twice at 65°C for 15 min under agitation in 0.1M NaHCO3 and 1% SDS as in^48^. After reverse crosslinking and DNA clean-up, 3 to 15 ng of immunoprecipitated DNA were used to prepare the sequencing libraries using the NEBNext Ultra II DNA Library Prep Kit (NEB, Ref: E7645S) according to the manufacturer’s instructions. Libraries were sequenced on a NovaSeq 6000 (Illumina) in SE100 mode.

### Whole exome sequencing

Genomic DNA from samples (DMSO-D0, DMSO-D147-#3, DMSO-D171-#5, DMSO-D131-#6, 5-FU-D67-#2, 5-FU-D153-#2, 5-FU-D50-#3, 5-FU-D147-#3, 5-FU-D171-#5 and 5-FU-D131-#6) were extracted with NucleoSpin Tissue, Mini kit for DNA from cells and tissue (Macherey Nagel, Ref: 740952.50) and sequenced on a NovaSeq 6000 (Illumina) with a 100X depth.

## Author contributions

JM, AD, CL, LB, SBT, AE and AT performed experiments. scChIP-seq experiments were conducted together with SF and KG. PDX experiments were performed by EM, LS and AD. MB and SB performed sequencing. PP and CV performed omics data analysis. Lineage barcoding data were analyzed by AML, CV and LP. Whole exome sequencing data were analyzed together with EL. CV, LP and JM conceived and designed experiments. CV, JM, PP and LP wrote the manuscript with input from all authors.

## Additional Information

Correspondence and requests for materials should be addressed to.

